# Experimental phasing of MicroED data using radiation damage

**DOI:** 10.1101/819086

**Authors:** Michael W. Martynowycz, Johan Hattne, Tamir Gonen

**Affiliations:** Howard Hughes Medical Institute, Departments of Biological Chemistry and Physiology, University of California Los Angeles, Los Angeles, CA 90095

## Abstract

We present an example of an experimentally phased structure using only MicroED. The structure of a seven-residue peptide is determined starting from differences to the diffraction intensities induced by structural changes due to radiation damage. The same wedge of reciprocal space was recorded twice by continuous rotation MicroED from a set of 11 individual crystals. The data from the first pass were merged to make a “low-dose dataset”. The data from the second pass were similarly merged to form a “damaged dataset.” Differences between these two datasets were used to calculate a Patterson difference map and to identify a single heavy atom site, from which initial phases were generated. The structure was then completed by iterative cycles of modeling and refinement.

## Introduction

Microcrystal electron diffraction (MicroED) is an electron cryo-microscopy (cryoEM) method that determines atomic resolution structures from vanishingly small three-dimensional crystals (Shi et al., 2013). The crystalline sample is exposed to the electron beam in diffraction mode under cryogenic conditions and the data is collected using a fast camera as a movie as the crystal is continuously rotating in a single direction (Nannenga et al., 2014). Continuous rotation MicroED has been used by several laboratories worldwide to determine structures of previously unknown proteins, protein complexes, ion channels, peptides, small molecules and inorganic material (Clabbers et al., 2017; de la Cruz et al., 2017; Gemmi et al., 2015; Gruene et al., 2018; Jones et al., 2018; Liu and Gonen, 2018; Palatinus et al., 2017, 2015; Rodriguez et al., 2015; Wang et al., 2018; Xu et al., 2018; Yonekura et al., 2015).

To date, all structures determined by MicroED were phased either by molecular replacement (MR) or *ab initio* direct methods. Direct methods are only applicable to crystals whose diffraction intensities extend to very high resolution; in practice to ∼1Å (Jones et al., 2018; Sawaya et al., 2016). Crystals that do not diffract to these resolutions can often be phased by molecular replacement if a homologous structure is available as a search model (McCoy et al., 2007; Scapin, 2013). In cases where no homologous structure is available and diffraction is worse than ∼1Å, phases must be recovered by other means. Phasing in electron crystallography of two-dimensional crystals has been previously demonstrated using images (Henderson et al., 1990; Henderson and Unwin, 1975) however to our knowledge no examples of successful experimental phasing have been reported using only MicroED data from three-dimensional crystals.

Experimental phasing methods attempt to solve the phase problem by calculating the phase of each structure factor from the diffraction intensities. Interpreting a density map in terms of an atomic model is the final step of crystallography analysis. To generate a density map, into which the atomic model is built, both the structure factor amplitudes and phases are required. Amplitudes are proportional to the square root of the reflection intensities, which are measured to sufficient accuracy by modern detectors, but the phase information is lost in MicroED measurements just like in X-ray crystallography giving rise to what is known as “the phase problem”(Rhodes, 1993; Taylor, 2003). The inability to directly measure the phase can be circumvented by using the phases from a homologous structure in molecular replacement (MR)(McCoy et al., 2007; Rossmann, 1990). For sufficiently small molecules, diffracting to very high resolution, phases can even be calculated *ab initio*(Hauptman, 1986; Sheldrick, 2007). For macromolecules without a sufficiently similar, known, structure, the phases must be determined experimentally. In X-ray crystallography, the anomalous scattering of heavy atoms results in measurable differences between the intensities of Friedel mates **I**_*hkl*_, and **I**_*-h-k-*_ *l*(Nakane et al., 2016). These differences allow for the determination of the heavy atom substructure of the protein using single or multiple anomalous dispersion (SAD, MAD)(Hendrickson, 1991; Rice et al., 2000). The heavy atom phases are then used to generate initial phases of the protein structure, and the resulting initial density maps may be generated. Electrons do not undergo this type of anomalous X-ray scattering(Parthasarathy, 1961) so this approach is not feasible with MicroED.

Before the widespread availability of tunable synchrotron sources necessary for SAD/MAD experiments, it was common to use heavy metal phasing approaches to solve the phase problem(Green et al., 1954; Perutz, 1956). Here the protein is crystallized and then resulting crystals were soaked in heavy atoms, such as lead or platinum. These heavy atoms scatter stronger than the lighter protein atoms. Collecting data from a crystal without the heavy atoms (the “native” dataset) and subtracting it from data taken from a derivative crystal with the heavy atom (the “derivative” dataset), would allow for the location of the heavy atoms and generation of initial protein phases. This technique is known as (single or multiple) isomorphous replacement (SIR/MIR) (De La Fortelle and Bricogne, 1997) because it requires that the lattice of the native and the derivative crystals are the same. However, soaking heavy atoms into proteins can often destroy the crystal or change the unit cell. Such non-isomorphism precludes calculating a meaningful difference map, and renders heavy atom location impossible, preventing structure determination altogether. Isomorphous replacement is effective in X-ray scattering because heavy metal atoms scatter much stronger than the light atoms typically found in biological materials. The difference in electron scattering between light and heavy atoms is much weaker in comparison, leading many to suggest that this approach would be intractable in electron diffraction measurements(Henderson, 1995).

Radiation damage from the X-ray beam has been used to generate initial phases without having to modify protein crystals (Ravelli et al., 2005, 2003; Thorn et al., 2012). This has been named radiation-induced phasing, or RIP (Nanao et al., 2005). Here, the exposure to the X-ray beam resulted in specific parts of the protein changing, such as the breaking of disulfide bonds(Galli et al., 2015; Ravelli et al., 2005). Partitioning the data into a damaged and undamaged set can locate these sites, using methods analogous to those used in anomalous dispersion or isomorphous difference experiments. This approach to generating initial phases is appealing, as it avoids the complication of crystallizing the sample under different conditions, soaking heavy metals into the delicate protein crystals, or accessing a tunable X-ray source. Non-isomorphism is also a much lesser concern since the two datasets are collected from the same crystal.

Here, we demonstrate that radiation damage from exposure to the electron beam can be used to solve the phase problem in MicroED experiments. Two sweeps of continuous rotation MicroED data were collected from 11 crystals of a polypeptide. Crystals of this peptide could only be grown in the presence of zinc acetate and it is known that a single zinc atom incorporates into asymmetric unit (J. Hattne et al., 2018). We denote the merged, first-pass data as the “low-dose” dataset. The merged data from the second pass we refer to as the “damaged” dataset. The difference of the amplitudes between these two datasets was calculated and used to identify the zinc site in the unit. The full atomic model was iteratively built out from the initial phases from this single heavy atom solution.

## Results

The diffraction images used to initially determine the structures with PDB IDs 6CLI and 6CLJ were integrated and truncated to 1.4Å resolution (J. Hattne et al., 2018). 6CLI corresponds to the structure of the peptide GSNQNNF after an average exposure of 0.17e^-^ Å^-2^, and 6CLJ corresponds to the same structure after an exposure of 0.5 e^-^ Å^-2^. While this structure was originally determined *ab initio* at a resolution of 1.0 Å, no direct methods solutions were possible with data at 1.4 Å. The structure factor amplitudes from the latter, damaged dataset were subtracted from the amplitudes from the former, low-dose dataset. Amplitudes for each reflection were used to calculate a difference dataset (**Figure 1**) and isomorphous difference Patterson maps were calculated for the difference dataset at 1.4Å resolution. The strongest peak in the difference Patterson was at the 3.6σ level(**Figure 1**). The other peaks present were all below the 3.5σ level. The strongest peak belonged to a cluster of vectors corresponding to distances between the zinc atom and others in the unit cell. This was confirmed by generating a list of all vectors from the atoms in the structure 6CLI. No vectors were present corresponding to the other discernable peaks of the difference Patterson map. Identification of heavy atoms in the difference dataset by Patterson methods found a single solution that corresponded to the zinc atom in this crystal structure. In this way, the location of the zinc site was found to be within 0.14Å from the previously refined location in 6CLI (J. Hattne et al., 2018). We similarly found very clear indications that the zinc atom was generating the difference signal by inspecting the difference Fourier map using the phases from 6CLI (**Figure 1, Figure 2**).

**Figure 1.**
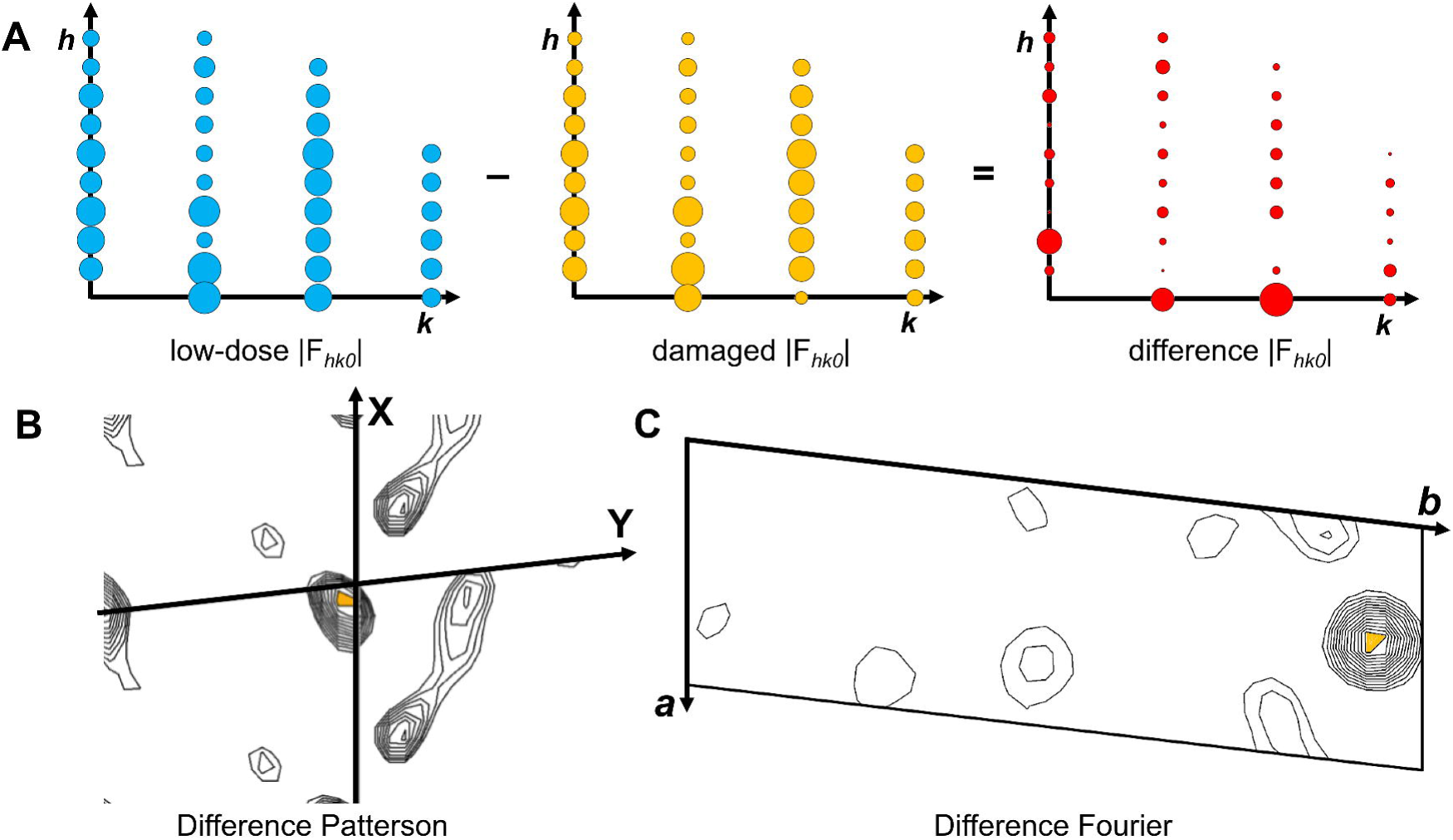
Differences between the low-dose and damaged datasets. (A) Reflection amplitudes for the upper quarter of the hk0 plane for the low-dose (blue), and damaged (yellow), and difference (red) datasets at 1.4Å resolution. Magnitude is indicated by the size of the circles. This is an approximate cartoon representation. (B) Isomorphous difference Patterson map viewed down the z/w axis at a resolution of 1.4Å with contours drawn every 0.2 *σ*. (C) Difference Fourier map viewed down the **c** axis with contours drawn every 1*σ*. The maximum peak of each map is colored in orange for clarity.

**Figure 2.**
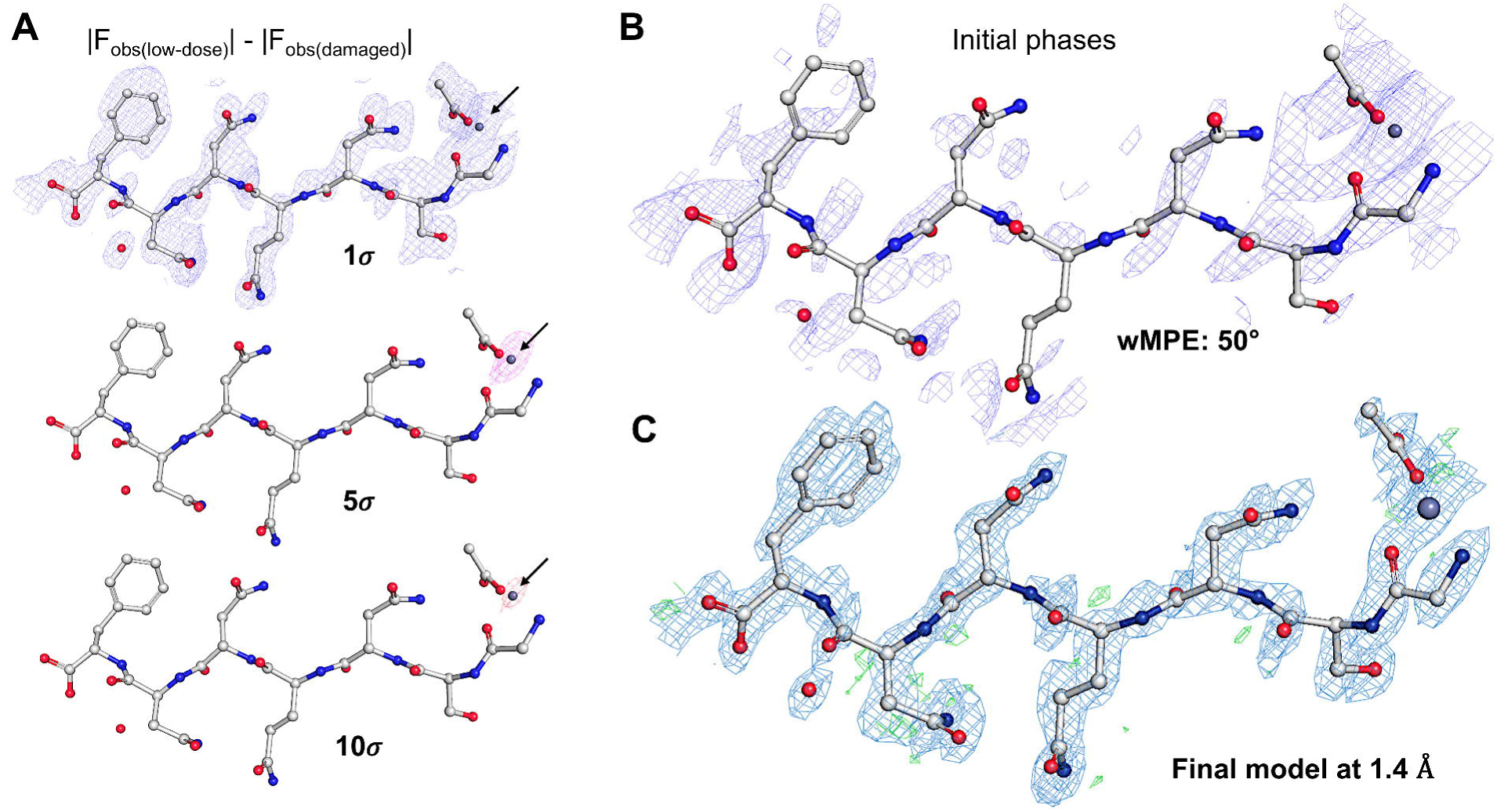
Locating the heavy atom substructure and solving the structure. (A) Observed difference maps between the damaged and undamaged structure of GSNQNNF using the phases of 6CLI at 1.4Å resolution. The map is contoured at 1*σ* (top), 5*σ* (middle), and 10*σ* (bottom) levels. Black arrows indicate the location of the zinc atom in the unit cell. (B) Density map at 1.0*σ* generated from the initial heavy atom solution. (C) The final structure solved at 1.4 Å resolution with the corresponding 2F_o_-F_c_ map in blue contoured at the 1.5*σ* level and F_o_-F_c_ map in green contoured at the 3.0*σ* level.

The heavy atom position was used to calculate phases, and an initial density map was generated (**Figure 2**). We calculated the accuracy of our solution using the weighted mean phase error (wMPE). This is a measure of the difference between the known phases of the final structure (6CLI), and those calculated during the solution process. Here, 0° would correspond to the perfect reproduction of the known phases, and 90° being completely random(Lunin and Woolfson, 1993). The initial solution from this single site corresponded to a wMPE of 50.5°. Structures in macromolecular X-ray crystallography are routinely determined from starting phases of similar quality(De La Fortelle and Bricogne, 1997; Nanao et al., 2005). Here, clear density was present around the zinc atom and in the surrounding areas of the structure (**Figure 2**).

Building the structure was done manually. Attempts at improving the density maps using automated density modification or phase improvement algorithms commonly used in X-ray crystallography were not successful. The initial maps from the single heavy atom site were poor so complete residues could not be built directly into such densities automatically. Building out the structure was instead initiated by manually adding single atoms into the initial density map and bootstrapping from there (**Figure 2**).

We initially built into the clear, individual peaks in the density. These corresponded to additional atom locations nearby the initial heavy-atom solution between symmetry-related heavy-atom sites. In each round, one to three carbon atoms were placed manually at full occupancy with atomic displacement parameters of 0. New maps were generated without further refining the atom positions. If the new maps did not improve the initial density as judged by the wMPE, these atoms were discarded and others were tested similarly. After 15 rounds of single-atom building, density for entire amino acids became visible allowing individual amino acids to be built into the density.

The final structure was completed in 34 rounds of manual building and modeling (**Figure 2, Table 1**). This solution was used to re-refine the damaged dataset at 1.4Å resolution to similar statistics (**Table 1**) as previously determined(Hattne et al., 2018). The all atom r.m.s.d between known and newly experimentally phased structure was 0.071Å.

**Table 1.** MicroED data from low-dose and damaged datasets.

|  |  | Low-dose<br>(0.17 e<sup>-</sup> Å<sup>-2</sup>) | Damaged<br>(0.5 e<sup>-</sup> Å<sup>-2</sup>) |
| --- | --- | --- | --- |
| Data analysis | Resolution range (Å) | 14.0 – 1.40<br>(1.51 - 1.40) | 13.97 - 1.40<br>(1.51 - 1.40) |
|  | Space group | P1 | P1 |
|  | Unit cell<br>(a,b,c)(Å)<br>(α,β,γ)(°) | 4.88, 14.17, 17.62<br>83.60, 84.98, 83.31 | 4.85, 14.15, 17.43<br>83.70, 85.43, 82.96 |
|  | Total reflections (#) | 6026 (1308) | 6202 (1445) |
|  | Multiplicity | 8.4 (8.5) | 8.7 (9.3) |
|  | Completeness (%) | 77.2 (78.5) | 78.1 (79.0) |
|  | Mean I/σ(I) | 8.5 (6.6) | 8.7 (6.7) |
|  | R<sub>meas</sub> | 0.22 (0.28) | 0.21(0.29) |
|  | CC<sub>1/2</sub> | 0.97 (0.94) | 0.98 (0.96) |
|  | R1 (%) | 22.2 | 24.3 |
|  | R<sub>complete</sub> (%) | 28.9 | 32.2 |

We chose to validate our structure using R_complete_ rather than the traditional R_free_ typically used in macromolecular crystallography(Brünger, 1992; Luebben and Gruene, 2015). Given the limited number of unique reflections (**Table 1**), assigning a fraction of them to a test set would make building an accurate model more difficult and not serve as an accurate validation of the structure. Although we have many observations with high redundancy in our data (**Table 1**), the total numbers of unique reflections for each of these structures were only ∼700 after merging at this resolution. For calculating R_complete_, the data from the final refinement was divided into 10 datasets, each withholding approximately 70-80 reflections for the free set. Each dataset was then refined using small perturbations to the initial model. Cross validations between the individual datasets gave the final R_complete_. This approach allowed us to refine and build the model using all the reflections. R_complete_ has been shown to accurately validate refinements without bias for both small molecule and macromolecular datasets(Clabbers et al., 2019).

## Discussion

We demonstrate that radiation damage can be used to generate initial experimental phases for MicroED data and that this information can be used for structure solution. The structure of GSNQNNF from these initial phases was determined at 1.4Å. At this resolution, our attempts at an *ab initio* direct methods solution were unsuccessful. This is consistent with past reports that demonstrated that resolutions of ∼1Å are required for direct methods solutions of MicroED and of X-ray data (Hauptman, 1986; Sawaya et al., 2016; Sheldrick, 2007). The initial phases in our analysis were determined by locating a single zinc atom that was particularly susceptible to radiation damage. The sensitivity of these atoms to damage in MicroED was noted in previous studies (Hattne et al., 2018). From these initial observations, we hypothesized that radiation damage would allow us to locate this zinc atom in a difference dataset calculated between the low dose dataset and the damaged dataset. Indeed, in real space, the difference showed a relatively strong signal for zinc (16*σ* level) while unmodeled amino acids contributed differences at the ∼2*σ* level (**Figure 2**).

Radiation-induced phasing is similar to an isomorphous replacement experiment using native and derivative datasets(Ravelli et al., 2005). The heavy atom, or derivative, dataset in our experiment corresponds to the low-dose dataset, and the native dataset corresponds to the damaged dataset because the zinc would have been ionized and removed from the lattice (Hattne et al., 2018; Nave, 1995). Rather than adding a heavy atom to a native dataset as is customary in isomorphous replacement, radiation damage reduces the contributions of the natively present atoms, such that it gradually disappears as the measurement progresses. Whereas the native dataset in an isomorphous replacement measurement has more scattering power than the derivative, the low-dose dataset in RIP scatters stronger than the that which has seen a higher exposure to the electron beam. In RIP, the difference between the atoms of these datasets manifests as a reduction of occupancy and/or an increase of B-factor in the damaged structure. Indeed, our final solutions demonstrated that the atomic displacement of the zinc in the low-dose dataset refined to a value of 4Å^2^, whereas it was 6Å^2^ in the damaged structure – both at full occupancy.

We initially demonstrated the location of the zinc atom from differences in the intensities of the low dose dataset and the damaged dataset. All attempts to improve the initial phases using density modification or phase improvement software failed. Only manually placing model atoms into the density resulted in improved wMPEs. The difficulty in improving the initial phases is to be expected, as the software has been developed using structures from X-ray crystallography, and has not been validated for MicroED experiments (Cowtan, 2010; Terwilliger, 2000).

This peptide only crystallized in the presence of a zinc atom that is present in the final model (Hattne et al., 2018). Although we did not perform a heavy metal soaking experiment *per se* our approach can be thought of as a combination of both RIP and SIR phasing: damage allowed us to locate a zinc site, and the fact that it was a heavy atom gave enough signal to begin building the model. Future studies may find that this also applies to charged residues that are also particularly susceptible to damage (Hattne et al., 2018). However, the phasing power from a small number of light-atom sites, such as oxygen, may not be sufficient to begin building an atomic model. We suspect that multiple sweeps of data can be collected, each with progressively more damage to the structure to find additional sites of specific damage. Examining the limits of RIP using MicroED data is an exciting avenue for future research and development.

We were able to demonstrate a structure solution from MicroED data using experimentally determined initial phases. It may now be possible to solve structures by collecting multiple datasets on the same crystals with different levels of exposure. The later datasets would have more damage manifested in the structure. This would change the integrated intensities, and initial phases may be generated from these differences. Here we could monitor the progress of phase improvement because the final structure of this peptide was known (Hattne et al., 2018). Picking the correct hand of an SIR/SAD solution typically relies on inspecting the maps from both hands after density modification (Wang, 1985). Because density modification procedures in electron diffraction are poor (Gonen et al., 2005; Wisedchaisri and Gonen, 2011) we were unable to accurately assess phase improvement from the initial map, it is required to build out both hands of the solution until a clear answer is determined. Nevertheless, we show that locating accurate positions of heavy atoms and building a correct model starting from experimentally determined initial phases is feasible using MicroED data. Future improvements to density modification and automatic building routines tailored to MicroED data may lead to automated structure solution pipelines that X-ray crystallographers currently enjoy.

## Acknowledgements

We would like to thank Michael R. Sawaya (UCLA) for helpful discussions. The data for this paper was originally collected by Calina Glynn (UCLA) at the Janelia Research Campus and published in Hattne et al., 2018. The Gonen laboratory is supported by funds from the Howard Hughes Medical Institute. Coordinates and maps for both damaged and undamaged structures have been deposited in the Protein Data Bank and the Electron Microscopy Data Bank under accession codes (XXXX, YYYY and AAAA, BBBB, respectively).

## Methods

MicroED data from PDB entries 6CLI and 6CLJ were integrated with XDS (Kabsch, 2010a) to a maximum resolution of 1.4Å (Hattne et al., 2018), scaled and merged in XSCALE (Kabsch, 2010b), and converted to SHELX format in XDSCONV(Kabsch, 2010b). Patterson maps were calculated using the isomorphous difference Patterson tool in the computational crystallography toolbox (cctbx)(Grosse-Kunstleve et al., 2002). Slicing of the Patterson and Fourier maps for visualization was done using MAPSLICER(Winn et al., 2011). Calculating the Patterson vectors from the known structure was done using VECTORS(Winn et al., 2011). Difference datasets were prepared in SHELXC (Sheldrick, 2010) with a DSCA factor of 0.85. The zinc site was identified using SHELXD(Schneider and Sheldrick, 2002). Generation of initial phases for the low-dose and damaged datasets were calculated in SHELXE without solvent flipping, the free lunch algorithm, or heavy atom refinement(Sheldrick, 2002). Model building was conducted in COOT (Emsley and Cowtan, 2004) and subsequent refinements of the models was performed using SHELXL (Sheldrick, 2015) using electron scattering factors (Peng, 1999). Fourier difference maps were generated in ANODE using the difference datasets from SHELXC at the specified resolutions and DSCA values (Thorn and Sheldrick, 2011). R_complete_ was calculated as described (Luebben and Gruene, 2015). All atom RMSD values were calculated in Pymol (Schrödinger LLC, 2014). Figures were generated in PYMOL(Schrödinger LLC, 2014) or Excel, and arranged in PowerPoint.

